# CauloBrowser 2.0: an AI-powered update

**DOI:** 10.64898/2026.11.05.764773

**Authors:** Berent Aldikacti, Vincent A. Lawal, Peter Chien, Jared M. Schrader

## Abstract

CauloBrowser is a systems-biology database for *Caulobacter crescentus* that organizes the largest repository of experimental data for this model bacterium. The original CauloBrowser provided a web interface for accessing these data, but the growing number of experimental datasets and advances in AI chatbots motivated an updated resource. We developed CauloBrowser 2.0 (CauloBrowser.org) as an open-source application that broadens access to a growing collection of published systems-level data. The CauloChat AI interface allows users to query these datasets in natural language, generate visualizations, and achieve complex biological insights. While the database is hosted freely on the web, users may also download the database and run it locally, reducing dependence on the web-hosted service and allowing laboratories to use compatible unpublished datasets. Altogether, CauloBrowser 2.0 provides a significant update for exploring diverse systems biology data in *C. crescentus*.

**Importance:** We still lack free, open databases to organize the plethora of systems-level data for model bacteria. CauloBrowser 2.0 organizes published systems-biology datasets for the bacterial model organism *Caulobacter crescentus*. This database provides an easily searchable platform for users to quickly examine data on a gene-by-gene basis. The database is also accessible via an AI chatbot interface, enabling rapid data retrieval for the user and supporting more complex queries across multiple data types. Altogether, CauloBrowser 2.0 helps lower the barrier to performing big data analyses to improve the rate of scientific discovery.

## Introduction

*Caulobacter crescentus* is a free-living aquatic Gram-negative α-proteobacterium that has served as a model for bacterial cell biology, asymmetric division, and cell-cycle regulation (1, 2). Its dimorphic life cycle produces two developmentally distinct daughter cells: a motile, non-replicative swarmer cell and a sessile, replication-competent, stalked cell. Because these transitions are highly reproducible and synchronized populations can be readily obtained, *C. crescentus* has been widely used to study how bacterial regulatory networks coordinate growth, polarity, differentiation, and cell-cycle progression (2–4). The completion of the *C. crescentus* genome sequence and subsequent use of genome-wide and time-resolved datasets revealed a coordinated genetic network controlling development (3, 5). As systems-level datasets have continued to accumulate, there is a need for an accessible database for gene-level, genome-wide, and systems-level interpretation (6).

The original CauloBrowser was released as a community systems-biology resource for *C. crescentus*, with a particular emphasis on datasets related to asymmetric cell division and cell-cycle regulation (6). It addressed a practical problem in the field: many high-throughput *Caulobacter* datasets were distributed across separate publications, supplementary files, and specialized formats. By bringing these datasets into a gene-centered interface, CauloBrowser served as the first dedicated *Caulobacter* database and helped researchers compare diverse datasets. Since the original release, the number of *Caulobacter* datasets has continued to expand. To support continued development, CauloBrowser 2.0 incorporates new datasets, a redesigned database that enables AI-chatbot queries, and support for local deployment.

## Results

CauloBrowser 2.0 presents curated *Caulobacter crescentus* systems biology data through four principal interfaces: <u>Gene Browser</u>, <u>Gene Viewer</u>, <u>CauloChat</u>, and <u>Download</u>. The resource integrates gene annotations, molecular summaries, and a growing collection of systems-level data from published studies (7–22) (Supplementary Table 1). The examples in Figures 1–4 use example gene queries to demonstrate how users navigate the interfaces, inspect available measurements, review experimental metadata, and move between different data displays. The same search and navigation workflow applies to any gene represented in CauloBrowser, using its gene name, NA1000 locus tag, or legacy CB15 locus tag.

**Table 1.**
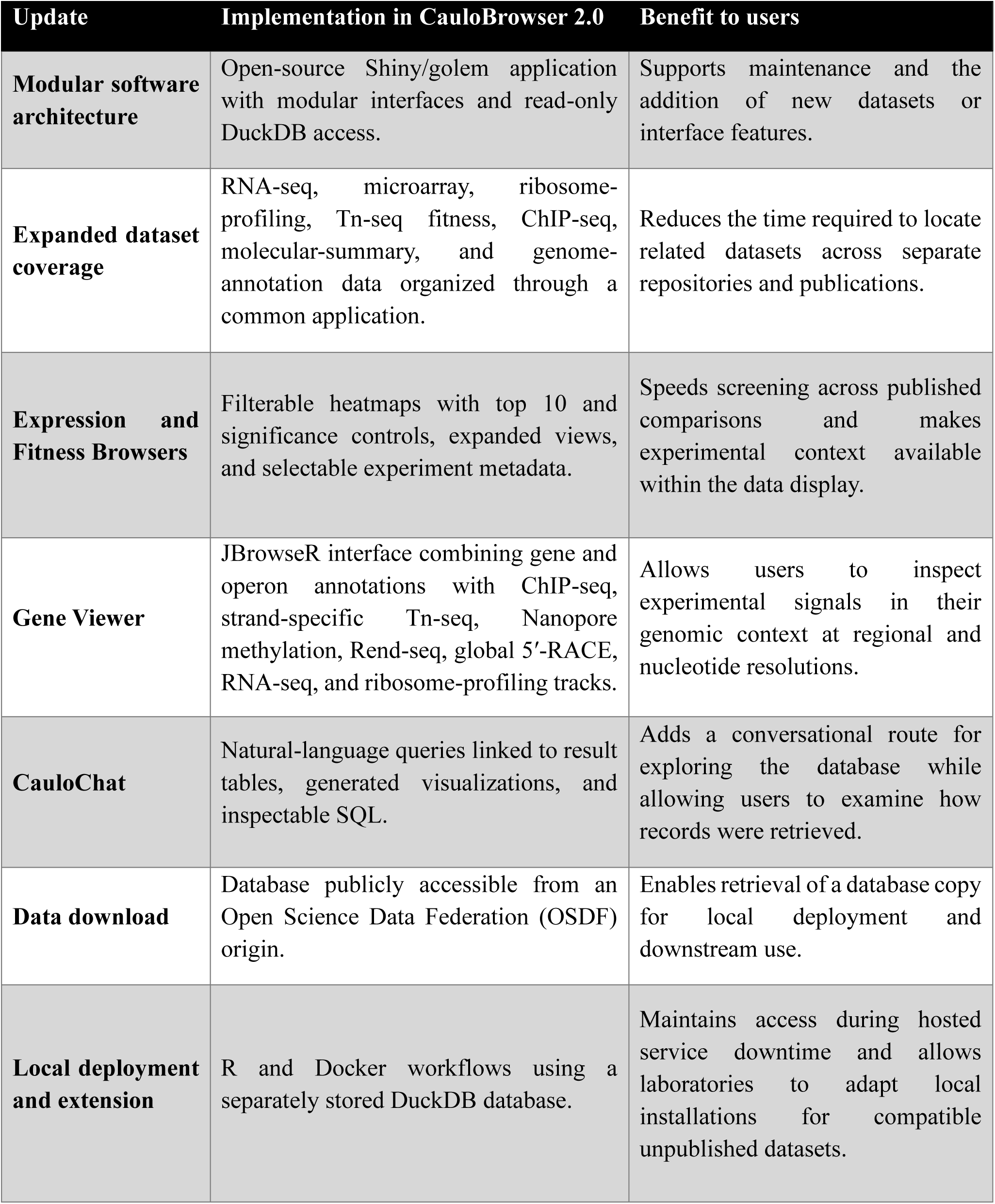
Major updates in CauloBrowser 2.0 and their benefits to users.

| **Update** | **Implementation in CauloBrowser 2.0** | **Benefit to users** |
|---|---|---|
| **Modular software architecture** | Open-source Shiny/golem application with modular interfaces and read-only DuckDB access. | Supports maintenance and the addition of new datasets or interface features. |
| **Expanded dataset coverage** | RNA-seq, microarray, ribosome-profiling, Tn-seq fitness, ChIP-seq, molecular-summary, and genome-annotation data organized through a common application. | Reduces the time required to locate related datasets across separate repositories and publications. |
| **Expression and Fitness Browsers** | Filterable heatmaps with top 10 and significance controls, expanded views, and selectable experiment metadata. | Speeds screening across published comparisons and makes experimental context available within the data display. |
| **Gene Viewer** | JBrowseR interface combining gene and operon annotations with ChIP-seq, strand-specific Tn-seq, Nanopore methylation, Rend-seq, global 5′-RACE, RNA-seq, and ribosome-profiling tracks. | Allows users to inspect experimental signals in their genomic context at regional and nucleotide resolutions. |
| **CauloChat** | Natural-language queries linked to result tables, generated visualizations, and inspectable SQL. | Adds a conversational route for exploring the database while allowing users to examine how records were retrieved. |
| **Data download** | Database publicly accessible from an Open Science Data Federation (OSDF) origin. | Enables retrieval of a database copy for local deployment and downstream use. |
| **Local deployment and extension** | R and Docker workflows using a separately stored DuckDB database. | Maintains access during hosted service downtime and allows laboratories to adapt local installations for compatible unpublished datasets. |

**Figure 1.**
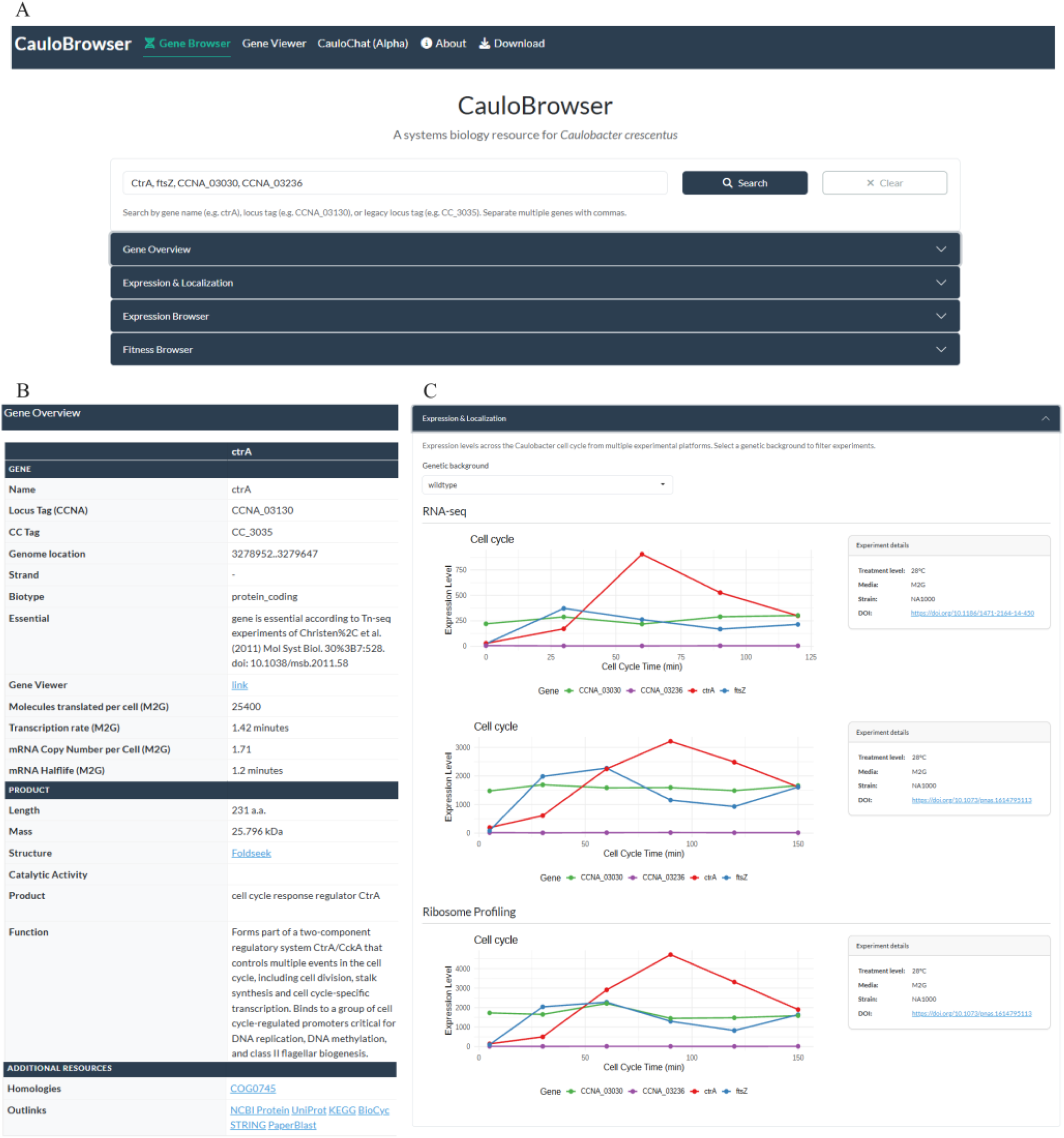
Gene Browser search, gene information, and cell-cycle data displays. (A) Gene Browser interface showing the search field populated with four example gene identifiers, the Search and Clear controls, and the expandable Gene Overview, Expression & Localization, Expression Browser, and Fitness Browser sections. The search field accepts gene names, NA1000 locus tags, and legacy CB15 locus tags. (B) Gene Overview table for a returned record. The displayed fields include gene and locus identifiers, genomic coordinates, strand, biotype, essentiality annotation, molecular measurements, product and functional descriptions, and links to the Gene Viewer and external resources. (C) Expression & Localization section filtered by genetic background. The interface displays two RNA-seq cell-cycle datasets and one ribosome-profiling cell-cycle dataset for the selected genes. Each plot is accompanied by its experimental conditions and source DOI, and consistent colors identify the genes across the three plots.

**Figure 2.**
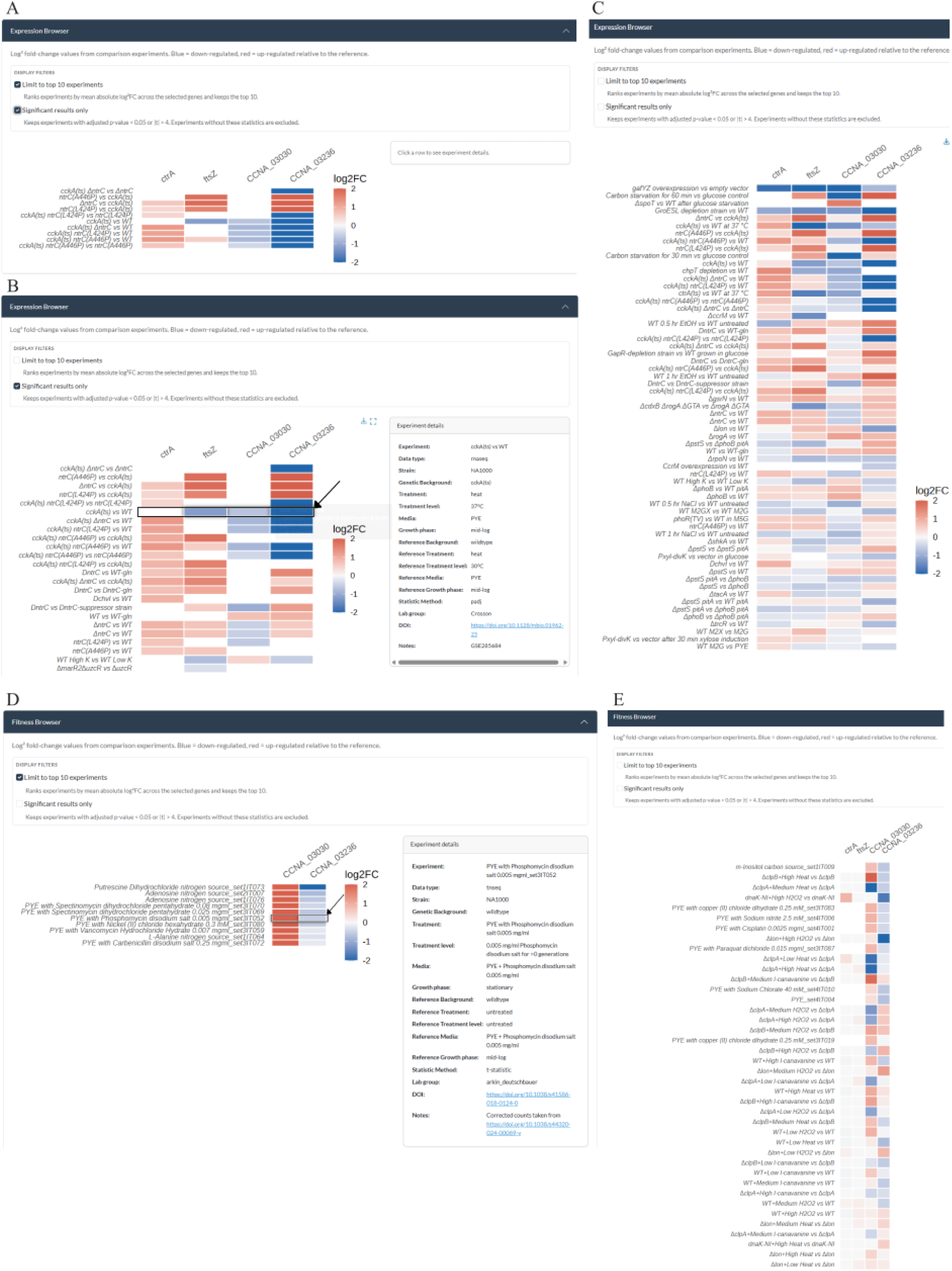
Expression and fitness comparisons within the Gene Browser. (A) Expression Browser heatmap with the top-10 and statistical-significance filters enabled. (B) Expression Browser with only the significance filter enabled and the details panel open for a selected comparison. (C) Expanded Expression Browser view with both display filters disabled. (D) Fitness Browser with the top-10 filter enabled, the significance filter disabled, and metadata displayed for a selected comparison. (E) Expanded Fitness Browser view with both filters disabled. In each heatmap, columns represent the queried genes and rows represent tested-versus-reference comparisons. Red and blue represent positive and negative displayed values relative to the reference, respectively, and color intensity indicates magnitude on the accompanying scale. The details panels provide the selected dataset’s tested and reference conditions, statistical method, source DOI, and associated notes.

**Figure 3.**
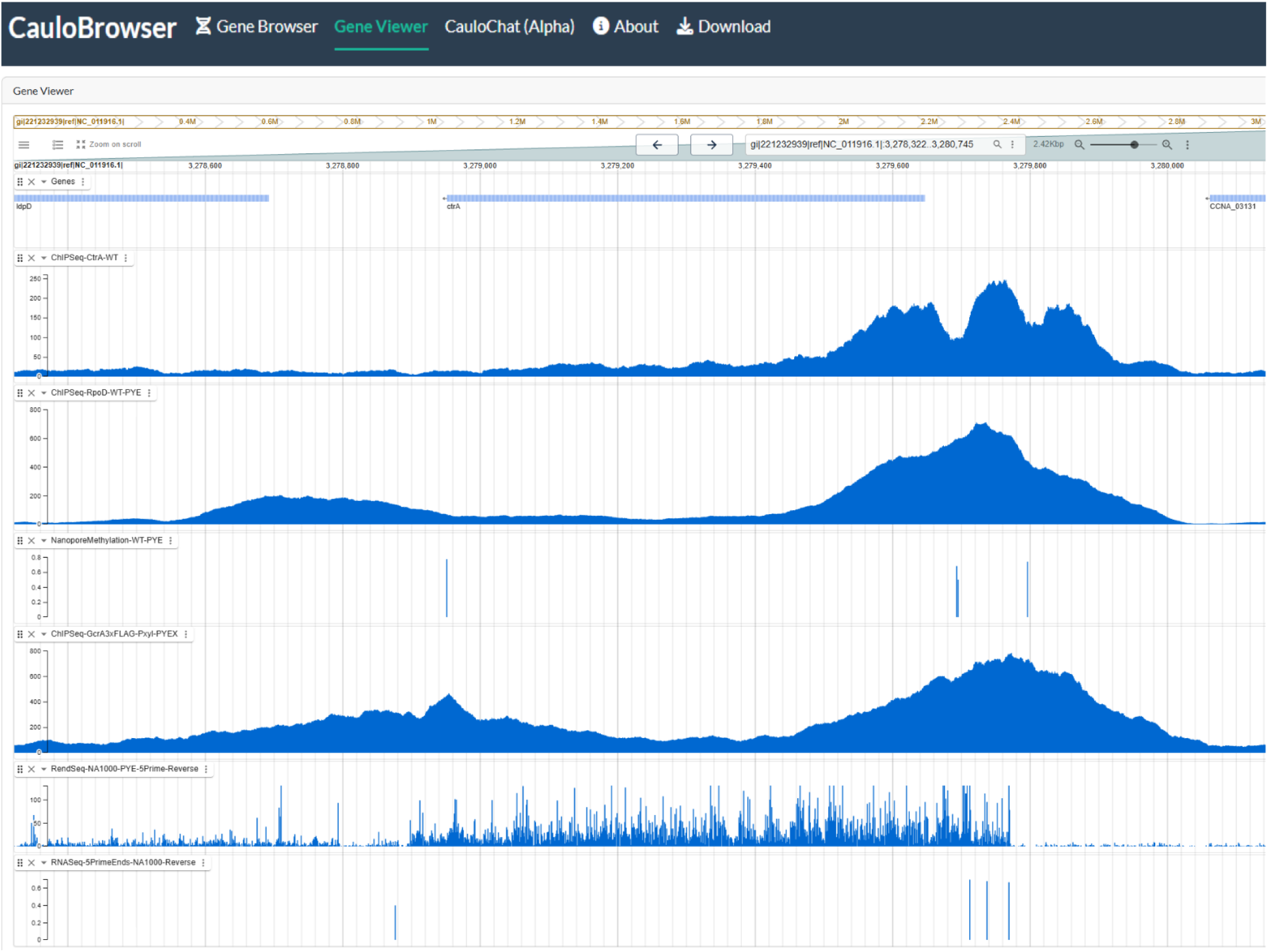
Gene Viewer displays genome and transcript annotations, as well as experimental signal tracks. Regional view surrounding the reverse-strand *ctrA* gene. Gene annotations are displayed together with CtrA, RpoD, and GcrA ChIP-seq profiles, Nanopore methylation, a reverse-strand Rend-seq 5′-end track, and a reverse-strand global 5′-RACE signal track used to map transcription start sites, all aligned on the same NA1000 genomic coordinate axis. Each quantitative track is displayed using its own vertical scale.

**Figure 4.**
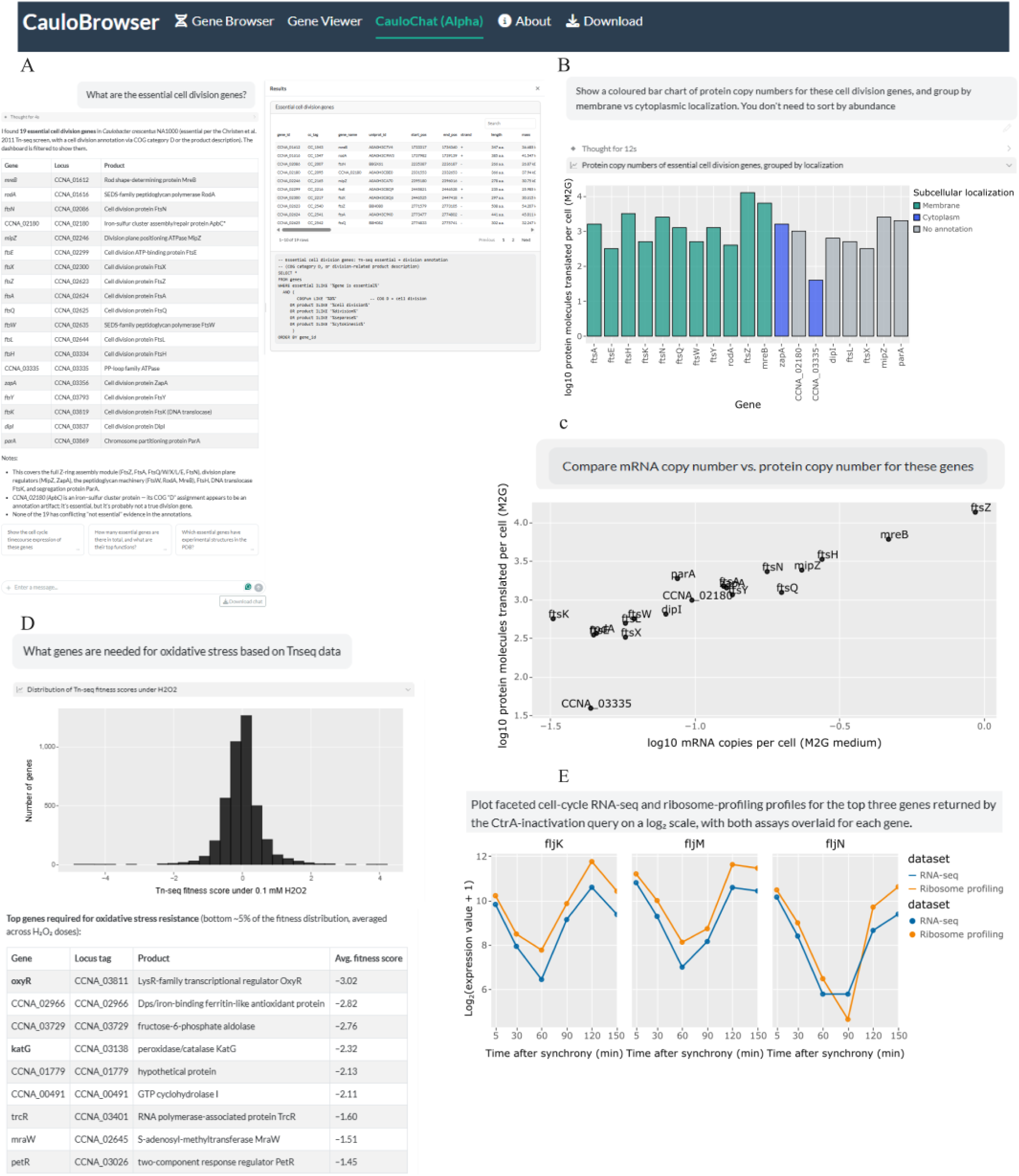
CauloChat supports natural-language database searches and quantitative visualization. (A) A request for essential cell-division genes returns 19 records selected by database annotations; the Results panel displays the records and the generated SQL. (B) A follow-up bar chart displays M2G estimates of protein molecules translated per cell for the selected genes, grouped and colored by subcellular-localization annotation. The vertical axis is log10-transformed; “no annotation” denotes missing localization information. (C) Further follow-up plots M2G mRNA-copy estimates against estimates of protein molecules translated per cell for the same genes on log10-transformed axes. (D) An oxidative-stress query displays a histogram of Tn-seq fitness scores from the 0.1 mM H₂O₂ comparison alongside a gene table summarizing scores across three H₂O₂-dose comparisons. (E) A query for genes responsive to CtrA inactivation applies an absolute log₂ fold-change threshold greater than 2 to the *ctrA(ts)*-versus-wild-type comparison and returns *fljK*, *fljM*, and *fljN* as the three highest-ranked records under the generated query. Faceted line-and-point plots compare their RNA-seq and ribosome-profiling measurements across six cell-cycle time points using log₂(expression value + 1).

### Gene Browser

The <u>Gene Browser</u> provides a common search interface for gene symbols, NA1000 locus tags, and legacy CB15 locus tags (Figure 1A). Users can enter one or several identifiers separated by commas and submit them in a single search. Results are organized into four expandable sections: <u>Gene Overview</u>, <u>Expression & Localization</u>, <u>Expression Browser</u>, and <u>Fitness Browser</u>. This organization allows the same set of search results to be examined across different information types without submitting a new query.

The <u>Gene Overview</u> displays the records returned for each queried identifier (Figure 1B). Available fields include the gene symbol, corresponding NA1000 and CB15 locus tags, genome coordinates, strand, biotype (protein-coding gene, non-coding RNA, etc.), essentiality annotation, product description, protein length and mass, absolute expression measurements in M2G medium, and links to external resources/databases. When several identifiers are submitted, the interface places their records in parallel columns so that users can compare the availability of annotations and measurements. Links from this section provide direct access to the corresponding <u>Gene Viewer</u> location and external resources, including NCBI Protein, UniProt, KEGG, BioCyc, STRING, PaperBLAST, Foldseek, and homology-group records.

The <u>Expression & Localization</u> section displays available cell-cycle measurements for the genes returned by the search (Figure 1C). A genetic-background selector filters the datasets shown in the panel. RNA-seq and ribosome-profiling datasets are presented as separate plots, with time on the horizontal axis and the measurement reported by the source dataset on the vertical axis. A consistent color is assigned to each queried gene across the plots. Each plot is accompanied by an experiment-details panel that identifies the strain, medium, temperature, and source DOI. The separate plots and metadata panels allow independently generated datasets to be viewed together while retaining their original sampling schedules, measurement scales, and experimental descriptions.

The <u>Expression Browser</u> and <u>Fitness Browser</u> extend the same search to differential-comparison datasets (Figure 2). In the <u>Expression Browser</u>, columns correspond to the queried genes and rows correspond to RNA-seq or microarray comparisons (Figure 2A–C). The heatmap displays the recorded log2 fold-change values, and the color key identifies positive and negative values relative to the reference condition. Users can apply the top-10 display option, apply the available significance criterion, or disable both options to view a broader set of comparisons. Selecting a row opens an experiment-details panel containing the tested and reference backgrounds, treatments, media, growth phases, statistical method, source DOI, and dataset notes. The <u>Fitness Browser</u> uses the same heatmap, filtering, and experiment-selection controls for transposon-fitness comparisons (Figure 2D and E). Maintaining a consistent interface between the <u>Expression Browser</u> and <u>Fitness Browser</u> reduces the need to learn separate controls for the two data types. The accompanying metadata identify the conditions and statistical information associated with each displayed comparison. Cells for which no value is displayed remain unfilled. Depending on the selected filters, an unfilled cell may represent an unavailable measurement or a result excluded by the current display settings; it should not be interpreted as a value of zero or as evidence of no effect.

### Gene Viewer

The <u>Gene Viewer</u> provides coordinate-based access to the NA1000 reference genome using JBrowseR (23) (Figure 3). It aligns the reference sequence, gene and operon annotations (24), and experimental signal tracks on a shared genomic coordinate axis. The regional view surrounding *ctrA* displays CtrA (15), RpoD, and GcrA (17) ChIP-seq profiles together with Nanopore methylation (19), a reverse-strand Rend-seq 5′-end track (20), and a reverse-strand global 5′-RACE signal track used to map transcription start sites (22). Displaying these datasets across the same genomic interval allows users to examine the locations of their signals relative to *ctrA* and neighboring annotated genes. Each quantitative track retains its own vertical scale, so their alignment provides positional context rather than a direct comparison of signal magnitudes.

Users can search for a coordinate or genomic feature, pan across the chromosome, and adjust the displayed interval. The Available tracks panel allows users to show or hide individual annotation and experimental tracks (Supplementary Figure 1A). Selecting the reverse-strand *ctrA* gene opens the Feature details panel, which displays its genomic position, length, strand, identifiers, gene biotype, available annotations, and feature sequence (Supplementary Figure 1B). The genome overview and navigation controls preserve the surrounding genomic context as users move between broader regional views and more focused inspection of individual features. Together, Figure 3 and Supplementary Figure 1 demonstrate how the Gene Viewer connects experimental signals with strand-aware genomic annotations across different spatial scales without requiring users to leave the application.

### CauloChat

<u>CauloChat</u> provides a natural-language interface to the CauloBrowser database (Figure 4). Users can request records, tables, or visualizations and refine their requests within the same conversation. Returned records are available in the Results panel, and expandable query details show the generated SQL. Users can inspect the query supporting a visualization and save the plot. A conversation history panel allows them to search for previous conversations or start a new one, and chat transcripts can be downloaded. CauloChat also provides explanatory text, which should be checked against the returned records and source studies for accuracy.

Figure 4A–C follows one conversation from a biological question to two views of the selected data. A request for essential cell-division genes returns 19 records matching the database’s essentiality and cell-division-related annotation criteria (Figure 4A). The response presents the genes with explanatory text, while the Results panel and generated SQL allow users to inspect the records and the criteria that selected them. Among the returned genes, CauloChat points to *ftsZ* in Z-ring assembly, *mipZ* in division-site positioning, and *ftsW* in cell-wall synthesis during division. This is a query-defined set, not a comprehensive catalogue of cell-division genes. Without restating the list, the user then requests a chart of estimates of protein molecules translated per cell, grouped by subcellular localization annotation (Figure 4B), followed by a plot of mRNA copies versus estimated protein molecules translated per cell for the same genes (Figure 4C). In its explanation of Figure 4B, CauloChat distinguishes nine membrane-annotated genes, three cytoplasm-annotated genes, and seven with no localization annotation, and notes that *ftsZ* has the highest displayed translation estimate, approximately 13,500 protein molecules translated per cell. The “no annotation” group indicates that localization information is missing. Its explanation of Figure 4C highlights differences among individual genes, including the contrast examined below. Together, these panels show how CauloChat carries a selected set of records into follow-up visualizations and provide both the displays and their explanations.

The measurements also illustrate how such exploration might prompt a question for further study. Within the selected set, *ftsZ* lies near the upper end of both the mRNA-copy and translation estimates, whereas *ftsK* has a low mRNA estimate but a moderate translation estimate (Figure 4C). The M2G mRNA-copy and translation estimates shown here do not, by themselves, establish differences in translation efficiency or protein stability. A testable hypothesis is that *ftsK* has relatively high protein output per transcript when measured in cultures grown under the same M2G conditions.

Figure 4D extends the examples from molecular-summary measurements to Tn-seq fitness data. In response to a question about oxidative stress, CauloChat displays the distribution of fitness scores from the 0.1 mM H₂O₂ comparison and a table of genes with low scores averaged across three H₂O₂-dose comparisons (10). Its accompanying explanation highlights *oxyR* (a central regulator of the H₂O₂ response) and *katG* (a catalase-peroxidase). This example shows how CauloChat helps users find relevant, data-based answers to their questions and identify results worth examining more closely. Although CauloChat retrieves well-known genes relevant to oxidative stress in this example, users are encouraged to double-check its explanations against the returned records and source study to guard against inaccurate AI-generated claims.

Another example shows how CauloChat can carry a gene selection from one experiment into a comparison of measurements from other datasets (Figure 4E). In response to a query about genes affected by CtrA inactivation, CauloChat uses the microarray comparison between a *ctrA(ts)* mutant and wild-type at 37 °C (25). The generated query applies an absolute log₂ fold-change threshold greater than 2 and returns *fljK*, *fljM*, and *fljN* as the three highest-ranked records. It then retrieves RNA-seq and ribosome-profiling measurements for these genes from the Schrader cell-cycle time courses (8) and displays both assays as faceted line-and-point plots across six time points on a shared log2(expression value + 1) scale. The profiles generally decrease from the early time points, reach minima at 60 or 90 min, and subsequently increase, with the RNA-seq and ribosome-profiling measurements showing broadly similar temporal patterns. This sequence demonstrates how CauloChat can connect a regulatory perturbation with focused exploration of cell-cycle measurements from other datasets. The response to CtrA inactivation does not establish that these genes are direct CtrA targets, and differences between the RNA-seq and ribosome-profiling curves do not by themselves measure translation efficiency.

Together, these examples show how CauloChat turns users’ natural-language queries into inspectable searches across CauloBrowser datasets, presenting records, visualizations, and explanatory text that users can evaluate to identify patterns and formulate testable hypotheses.

### Download

CauloBrowser can be installed and run locally from its open-source repository (Figure 5). The documented workflow provides two installation routes. For an R installation, users install the package from GitHub, retrieve the DuckDB database via Pelican, set the database location with CAULOBROWSER_DB_PATH, and launch the application. For containerized deployment, users install Docker, retrieve the DuckDB database, mount it when starting the CauloBrowser container, expose port 3838, and open the application in a web browser. Both routes allow laboratories to operate CauloBrowser independently of the hosted service while retaining a local copy of the database used by the installation. Detailed installation and local-deployment instructions are available in the CauloBrowser GitHub repository (https://github.com/baldikacti/caulobrowser).

**Figure 5.**
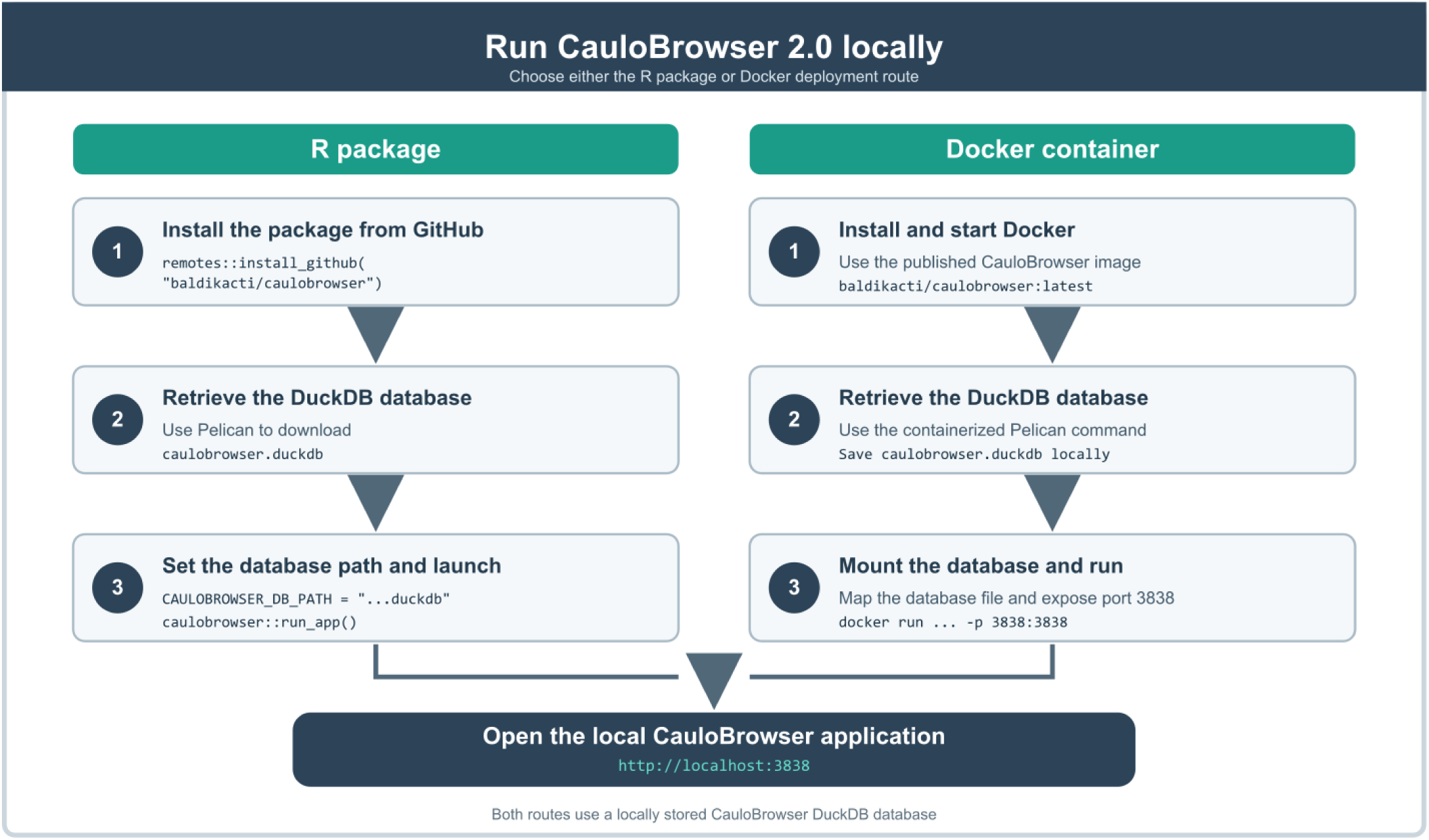
Full database download for local deployment of CauloBrowser 2.0. Local-deployment workflow showing the supported R package and Docker routes. In the R route, users install the CauloBrowser package from GitHub, retrieve the DuckDB database using Pelican, specify the local database path, and launch the application. In the Docker route, users obtain the published container image, retrieve the database with the containerized Pelican command, mount the database file, and expose port 3838 when starting the container. Both routes use a locally stored CauloBrowser DuckDB database and provide access to the application through http://localhost:3838.

## Discussion

CauloBrowser 2.0 extends the original resource by integrating gene-centered and genome-centered views of published *Caulobacter crescentus* systems biology data. An integrated framework allows users to move among annotations, cell-cycle measurements, differential-expression results, fitness data, and genomic context without repeatedly reconciling gene records across separate sources. This integration supports dataset discovery and contextual comparison while preserving distinctions among assays. Because RNA abundance, translation, mutant fitness, essentiality, and genome-track signals are not interchangeable, CauloBrowser retains assay-specific units and displays scales together with experimental conditions, statistical information, dataset descriptions, and source links. Users can therefore screen related datasets within a single interface and consult the original studies for methodological details and biological interpretation.

CauloChat extends access to the resource by allowing users to begin with a natural-language question rather than a specific database field or identifier. As Figure 4 illustrates, users can carry a selected gene set into follow-up plots, examine fitness data for a condition, or use genes selected from one experiment to explore measurements from another, then decide which patterns warrant a testable follow-up question. The Results panel and expandable query details provide an important level of transparency by allowing users to inspect the records returned and the database operation used to assemble the response. This is particularly relevant for a large-language-model interface because the scope of a response depends on the wording of the question, the available database fields, and the criteria generated from them. A query-defined group of genes may therefore differ from a manually curated biological category. CauloChat is presented as an alpha feature (indicating it is still under active development), and its responses should be treated as an entry point for exploration. Users should examine the returned records, the generated query, the experimental metadata, and the cited source studies before using an answer to support a biological conclusion.

CauloBrowser 2.0 also changes the software architecture and distribution of the original resource (6). The application is implemented as an open-source R package, uses a separately distributed DuckDB database, and can be run either in R or with Docker. Local deployment allows laboratories to maintain a database copy, associate it with a specific application version, and continue using the resource when the hosted service is unavailable due to scheduled downtime or institutional information technology shutdowns. Access to the source code and database schema also allows laboratories to adapt a local installation for their own unpublished datasets. After the measurements and associated metadata are organized according to the CauloBrowser database structure, they can be explored through the same interfaces while remaining within the laboratory’s local environment.

The usefulness of the resource remains dependent on the coverage and quality of the data it incorporates. Experimental metadata and statistical information are not reported uniformly across published studies, and some genes or comparisons consequently have more complete records than others. CauloBrowser preserves these differences rather than inferring missing information, and it does not replace assay-specific reanalysis or consultation of the original publications. As the public database expands, new records will be paired with source identifiers, tested and reference conditions, measurement definitions, statistical fields, and database-release information to preserve their interpretation and traceability. Continued evaluation of CauloChat will accompany the expansion of its query tools and supported question types. With these qualifications, CauloBrowser 2.0 provides a maintainable framework for connecting gene annotation, expression, translation, fitness, regulatory tracks, and genome context in C. crescentus systems biology.

## Materials and methods

### Application architecture and database organization

CauloBrowser 2.0 is implemented in Shiny for R using the golem framework. At application startup, the database path is obtained from the CAULOBROWSER_DB_PATH environment variable, and the supplied DuckDB database is opened in read-only mode. The connection is checked for six required tables: genes, experiments, experiment_conditions, de_results, timecourse_expression, and gene_viewer_metadata. This structure separates gene annotations, dataset metadata, experimental conditions, differential-comparison results, time-course measurements, and Gene Viewer configuration.

One row in the experiments table represents either one tested-versus-reference comparison or one complete time-course dataset; it does not represent an individual sample or biological replicate.

### Gene Browser data curation and implementation

The Gene Browser draws on 349 experimental datasets: 68 RNA-seq differential comparisons, two RNA-seq time-course datasets, 35 microarray differential comparisons, 243 Tn-seq fitness comparisons, and one ribosome-profiling time-course dataset (Supplementary Table 1). The microarray collection comprises 28 NA1000 comparisons and seven CB15 comparisons. The Tn-seq collection comprises 198 comparisons from the broad bacterial mutant-fitness study of Price *et al.* (9) and 45 comparisons from the stress-focused study of Aldikacti *et al.* (10). The Gene Browser also provides three gene-level molecular-summary datasets: mRNA copies per cell, mRNA half-life, and molecules translated per cell (11, 12).

Dataset selection, numerical data retrieval, and metadata curation were performed manually. Available gene-level data tables were downloaded from NCBI GEO or from supplementary files associated with the source publication. Candidate RNA-seq datasets were identified by searching NCBI GEO for *Caulobacter crescentus* studies, with particular attention to records using the NA1000 strain. Available processed fitness tables were downloaded manually from the study repositories or supplementary materials and checked against the corresponding publications. Microarray result tables were obtained from GEO when deposited there or from supplementary files associated with the source publication. Seven microarray datasets do not have a GEO accession in the supplied metadata and are therefore linked through their source publications. Experimental metadata were obtained from the repository record, the corresponding publication, or both. For each dataset, the strain, genetic background, treatment, treatment level, growth phase, medium, corresponding reference conditions, statistical method, laboratory, DOI, repository accession, and curation date were recorded when available. Gene-level values were converted to the database structure and linked to the corresponding dataset and gene records through experiment_id and gene_id.

A parameterized database query compares each term with the gene symbol, NA1000 locus tag, and legacy CB15 locus tag fields. Returned records are ordered according to the sequence of the submitted search terms. Gene records were standardized around NA1000 CCNA locus tags, while gene symbols and legacy CB15 CC locus tags were retained as alternative search identifiers. Product descriptions, functional annotations, essentiality information, genomic coordinates, and external database identifiers were stored with the corresponding gene when available. Essentiality annotations were obtained from the transposon-based analysis of Christen *et al.* (13). The M2G molecular summaries were mapped to NA1000 gene records from their source studies. These fields include mRNA copies per cell (copynumber), molecules translated per cell (translated_prot_ave), and mRNA half-life (mrna_halflife) (11, 12). Translation values are treated as estimates of protein synthesis derived from ribosome profiling rather than as direct measurements of steady-state protein abundance (11).

The time-course datasets comprise the Fang *et al.* RNA-seq series (7; GSE46915), the Schrader et al. RNA-seq series (8; GSE68200), and the Schrader et al. ribosome-profiling series (8; GSE68200). Available gene-level measurements and sampling information were downloaded from the associated GEO records and checked against the corresponding publications. Gene identifiers were mapped to NA1000 locus tags, and the measurements were converted to one record per gene, dataset, and sampled condition. Time-point labels, their plotted order, measurement values, strain, genetic background, medium, temperature, and source DOI were retained from the source records. The time-course view joins timecourse_expression with the gene, dataset, and condition tables and filters the returned records by the selected genetic background. The Fang series is displayed at 0, 30, 60, 90, and 120 min, whereas the Schrader RNA-seq and ribosome-profiling series are displayed at 5, 30, 60, 90, 120, and 150 min. Each dataset is plotted separately using its source-reported values and condition metadata. Lines connect the displayed observations, and a consistent gene-color assignment is used across panels. Values are not pooled or renormalized across studies or assay types.

For display, gene-level values are retrieved from de_results and joined to the corresponding gene and dataset metadata through gene_id and experiment_id. RNA-seq, microarray, and Tn-seq datasets are distinguished by the data_type field. A diverging scale centered on zero displays negative values in blue and positive values in red. For RNA-seq and microarray comparisons, the sign indicates the direction of expression relative to the reference. For Tn-seq comparisons, a positive value indicates relative enrichment of mutants disrupted at that locus, whereas a negative value indicates relative depletion in the source comparison (9, 10).

The significance option retains results with an adjusted *P value* below 0.05 when stat_method is recorded as padj, or an absolute *t statistic* greater than 4 when stat_method is recorded as t-statistic. Results without the applicable statistical value do not pass this filter. The microarray metadata supplied for the current inventory do not specify a compatible statistical method. Consequently, these datasets are available when the significance filter is disabled but are not retained by the implemented significance filter. The top-10 option ranks comparisons according to the largest absolute value recorded for any of the queried genes and retains up to ten comparisons. A nonblank heatmap cell represents a displayed result. A blank cell may indicate that a measurement is unavailable or that it was excluded by an active filter; it is not interpreted as a value of zero or as evidence of no expression response, fitness effect, or biological function.

### Gene Viewer configuration and track preparation

The Gene Viewer is implemented with JBrowseR (23) using the NC_011916.1 reference sequence. Its configured collection includes gene and operon annotations together with experimental tracks from ChIP-seq (15–18), plus- and minus-strand Tn-seq insertion tracks (10), Rend-seq 5’-end data (20), RNA-seq and ribosome-profiling (8), global 5′-RACE data used to map transcription start sites (22), and Nanopore methylation datasets (19). The application reads the reference assembly, searchable sequence index, feature tracks, and experimental tracks from the gene_viewer_metadata table. Gene and operon annotations are stored as indexed GFF or GFF3 feature tracks, whereas experimental signals are provided in browser-compatible indexed formats, including BigWig where appropriate. Each configured track is registered with a display name, track type, and network-accessible storage path. Available track files were assembled manually and converted to browser-compatible formats where necessary. The operon feature track is based on the transcriptome-derived annotation of Bharmal *et al.* (24).

Genes and operons are displayed with their genomic positions and orientations, and each quantitative signal track retains its own vertical scale. Selecting an annotated feature opens the Feature details panel, which reports the feature’s genomic coordinates and stored attributes and provides its sequence. The displayed tracks support coordinate-based inspection. CauloBrowser does not perform additional peak calling, cross-track normalization, binding-site prediction, or transposon-insertion analysis.

### CauloChat implementation and query-to-visualization workflow

CauloChat is implemented as a chatbot using querychat (26) and ellmer (27) and uses a read-only connection to the CauloBrowser database. The public implementation configures an OpenAI-compatible client using the Qwen3.8-27B model (28) through the Harmony HPC LiteLLM endpoint and enables filter, query, and visualization tools.

A natural-language request is interpreted using the available database fields and translated into a database query. The generated SQL is executed against the read-only database, and the returned records can be presented as explanatory text, a table, or a visualization. Follow-up requests remain within the same conversation, allowing records returned by an earlier query to be used in subsequent queries or visualizations without requiring the user to restate the complete request. Visualizations are generated from the measurements returned by the database query. Axis titles and transformations identify the fields and scales used in the resulting display. Users can inspect the generated SQL, save the resulting plot, and download the conversation as a Markdown transcript. CauloChat does not alter the curated records.

The selection criteria, data sources, and plotted transformations in the examples in Figure 4 were documented from the generated SQL, the accompanying results tables, and the downloaded CauloChat conversation transcripts. CauloChat selected the cell-division gene set using the database’s essentiality annotation, along with cell-division-related functional or product terms and then displayed the M2G molecular-summary measurements for those records. The oxidative-stress example queried Tn-seq comparisons of wild-type cells exposed to H₂O₂; its histogram used the 0.1 mM comparison, whereas the accompanying gene summary reported scores averaged across three H₂O₂-dose comparisons.

For the CtrA example, the generated query filtered the *ctrA(ts)*-versus-wild-type microarray comparison at 37 °C (25) for records with an absolute log₂ fold change greater than 2, ranked the returned records by absolute log₂ fold change, and retained the three highest-ranked records: *fljK*, *fljM*, and *fljN*. The query then retrieved RNA-seq and ribosome-profiling measurements for these genes from the Schrader cell-cycle time courses (8). Values from the six time points were transformed as log₂(expression value + 1) and displayed as faceted line-and-point plots; no within-gene or assay-specific rescaling was applied. These selections and transformations document the generated query and do not constitute an independent reanalysis of the source experiments.

### Data download and local deployment

The application can be installed from GitHub as an R package or run using the documented Docker workflow. The database is distributed separately, and its location is supplied through CAULOBROWSER_DB_PATH for native R use or mounted into the container for Docker use. The repository provides instructions for retrieving the database with Pelican. For containerized deployment, port 3838 is exposed to the host system, and the application is accessed through a local web browser. The application opens the supplied database in read-only mode.

Laboratories can prepare a local DuckDB database that follows the required table structure and use CauloBrowser with compatible unpublished datasets. Dataset preparation and conversion occur outside the running application, after which the local database path is supplied through the same configuration used for the distributed database. This allows a locally maintained data snapshot to be paired with a particular version of the application. The public code is distributed under the MIT license.

## Data availability

CauloBrowser 2.0 source code and installation instructions are available at https://github.com/baldikacti/caulobrowser, and the hosted application is available at https://caulobrowser.org. Supplementary Table 1 provides the supplied dataset inventory. Source publication and accession information accompany the corresponding data views. The repository documents retrieval of the separately distributed DuckDB database. Users should cite the original CauloBrowser publication, this update, and the source datasets used in their analyses.

## Acknowledgments

We thank members of the Schrader and Chien laboratories for testing CauloBrowser 2.0 and providing feedback during its development. CauloBrowser 2.0 is hosted on Harmony, a Kubernetes-based computing infrastructure developed with support from the National Science Foundation’s Campus Cyberinfrastructure (CC*) program (2430001). This work was supported by Indiana University startup funds to J.M.S., University of Massachusetts internal funds, and the Models to Medicine Center of the Institute for Applied Life Sciences at the University of Massachusetts.

## Author contributions

B.A. and V.L. contributed equally to this work. B.A. led software implementation, deployment infrastructure, and application development. V.L. contributed to data curation, manuscript preparation, project evaluation, and documentation. J.M.S. and P.C. supervised the project, provided biological and systems-biology guidance, and contributed to manuscript development.

## Competing interests

The authors declare no competing interests.

## Supplementary figures and figure legends

**Supplementary Figure 1.**
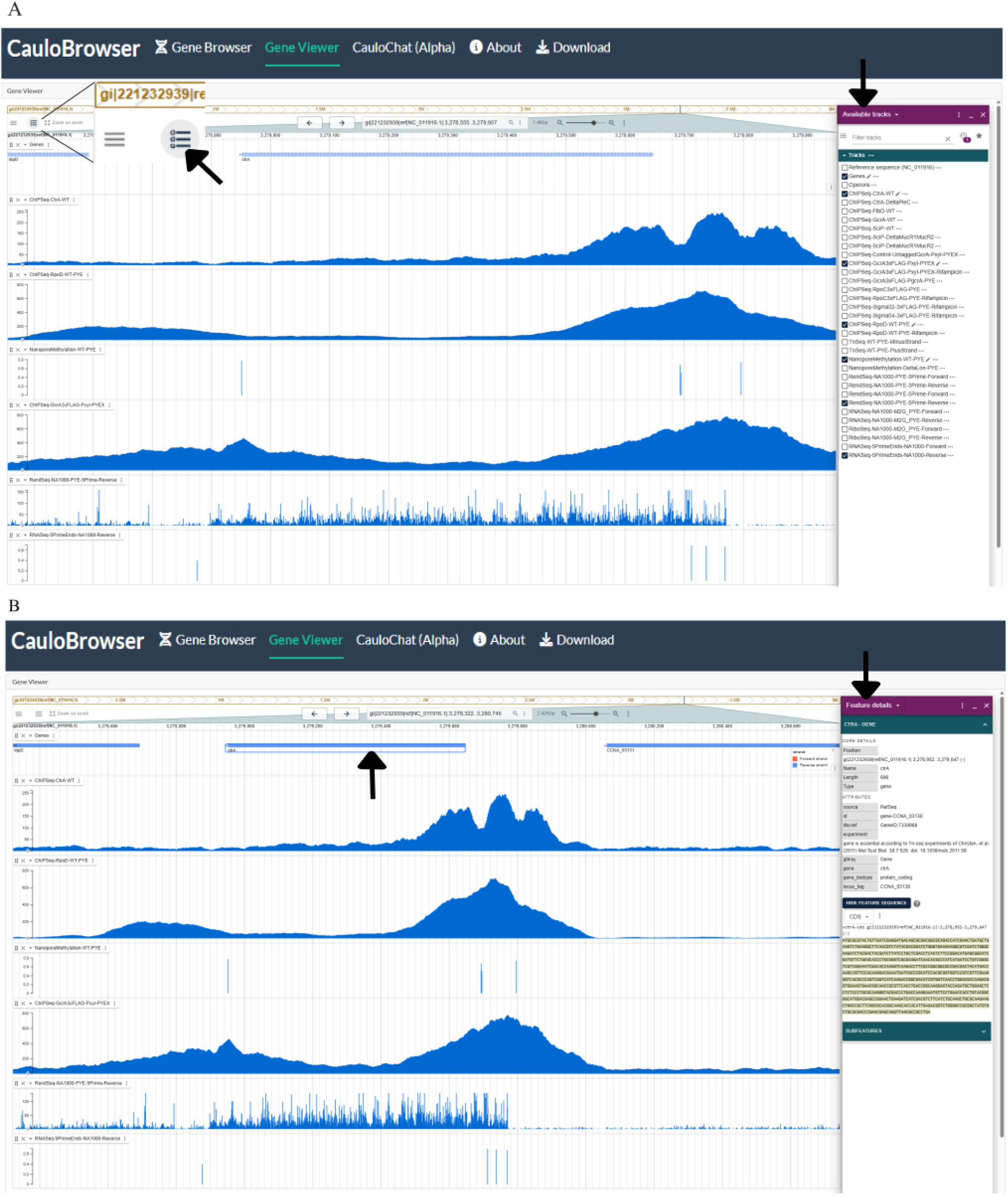
Gene Viewer track selection and feature inspection. (A) Regional Gene Viewer display with the Available tracks panel open. The arrows indicate the track-selection control and the panel used to show or hide reference, annotation, and experimental tracks. (B) Selection of the reverse-strand *ctrA* gene and the resulting Feature details panel. The arrows indicate the selected gene feature and the panel displaying its genomic position, length, strand, identifiers, gene biotype, stored annotations, and feature sequence.

